# Macrophage migrates on alternate non-adhesive surfaces

**DOI:** 10.1101/2022.08.10.503454

**Authors:** Fulin Xing, Hao Dong, Jianyu Yang, Chunhui Fan, Mengdi Hou, Ping Zhang, Fen Hu, Jun Zhou, Liangyi Chen, Leiting Pan, Jingjun Xu

**Affiliations:** The Key Laboratory of Weak-Light Nonlinear Photonics of Education Ministry, School of Physics and TEDA Institute of Applied Physics, Nankai University, Tianjin 300071, China; State Key Laboratory of Medicinal Chemical Biology, Frontiers Science Center for Cell Responses, College of Life Sciences, Nankai University, Tianjin 300071, China; State Key Laboratory of Membrane Biology, Institute of Molecular Medicine, National Biomedical Imaging Center, Center for Life Sciences, School of Future Technology, Peking University, Beijing 100871, China; Shenzhen Research Institute, Nankai University, Shenzhen 518083, China

**Keywords:** Macrophage, Cell-patterning, Cell migration, Non-adhesive surface, Podosome, Myosin IIA

## Abstract

Macrophages migrate across tissues upon immune demand, but their motility on heterogeneous substrates remains unclear. Protein-repelling reagents, *e*.*g*., poly(ethylene) glycol (PEG), are routinely employed to resist cell adhering and migrating. Contrary to this perception, we discovered a unique locomotion of macrophages *in vitro* that they overcome non-adhesive PEG gaps to reach adhesive regions in a mesenchymal mode. Adhesion to adhesive regions was a prerequisite for macrophages to perform further locomotion on the PEG regions, or else they kept a suspended round shape. Podosomes were found highly enriched on the PEG region, which supported macrophage migration. Myosin IIA played a negative role in macrophage motility. Moreover, a developed cellular Potts model reproduced the experimental observations. These findings uncovered a new migratory behavior on non-adhesive surfaces in macrophages.

**One-Sentence Summary:** Macrophages can migrate across non-adhesive surfaces that are absolute boundaries for other cell types.

---

**M**acrophages, serving as the first line of defense against exogenous pathogen, play crucial roles in various physiological and pathological processes *(1-4)*. Tissue-resident macrophages need to patrol through complex interstitial tissues to the injured or inflamed sites *(5,6)*. On the other hand, circulating monocytes perform transendothelial migration through tissues and differentiate into mature macrophages to execute their specific functions *(7,8)*. Therefore, the motility of macrophages is of great importance for immune surveillance.

Single-cell migration involves two distinct modes, known as the adhesion-dependent mesenchymal mode, and adhesion-independent but contraction-driven amoeboid mode *(9)*. Macrophages usually displayed a classic mesenchymal mode of migration in heterogeneous interstitial tissues *in vivo (10,11)*, with a relatively low speed compared to fast amoeboid cells, *e*.*g*., neutrophils. They also exhibit plasticity in morphology and migration, enabling them to deal with complex substrates, such as amoeboid migration on soft substrates or 3D environments and mesenchymal migration on stiff substrates *(12-14)*. Extensive research has focused on the migration properties of macrophages on homogeneous substrates. In the present work, based on a photolithography cell-patterning technique, we investigate the motility of macrophages in heterogeneous substrates with alternate non-adhesive gaps.

## Macrophages can migrate across alternate non-adhesive surfaces in a mesenchymal mode

To fabricate a heterogeneous substrate, we used an alternate non-adhesive surface composed of poly-L-lysine-g-PEG (pLL-PEG, abbreviated as PEG in the following sections) and fibronectin (FN) (Fig. S1A) with stripe and triangular lattice circular patterns (Fig. S1B). Protein-repelling regents such as PEG prevent cell adhesion via resisting non-specific protein adsorption *(15)*. Challenging this popular perception, we found that macrophages could actively migrate across the non-adhesive PEG region (Fig. 1A; movie S1, part I). The cells exhibited no directional migration, but random locomotion, as visualized by the irregularity of the migration trajectory (Fig. 1A). Unexpectedly, macrophages migrated over the non-adhesive region in a classical mesenchymal mode, other than an amoeboid morphology (Fig. 1B; movie S1, part II). In detail, the cell first extended out of the FN region with a subsequent elongation of the cell body. Afterwards, the leading edge reached another FN region, followed by the uropod dissociation and contraction of the cell body. The forward velocity of the leading edge on PEG surface was 0.68±0.04 μm/min, which was lower than that on the glass surface (0.92±0.05 μm/min) (Fig.1, C and D), indicating that PEG surface was not conducive for cell locomotion.

**Figure 1.**
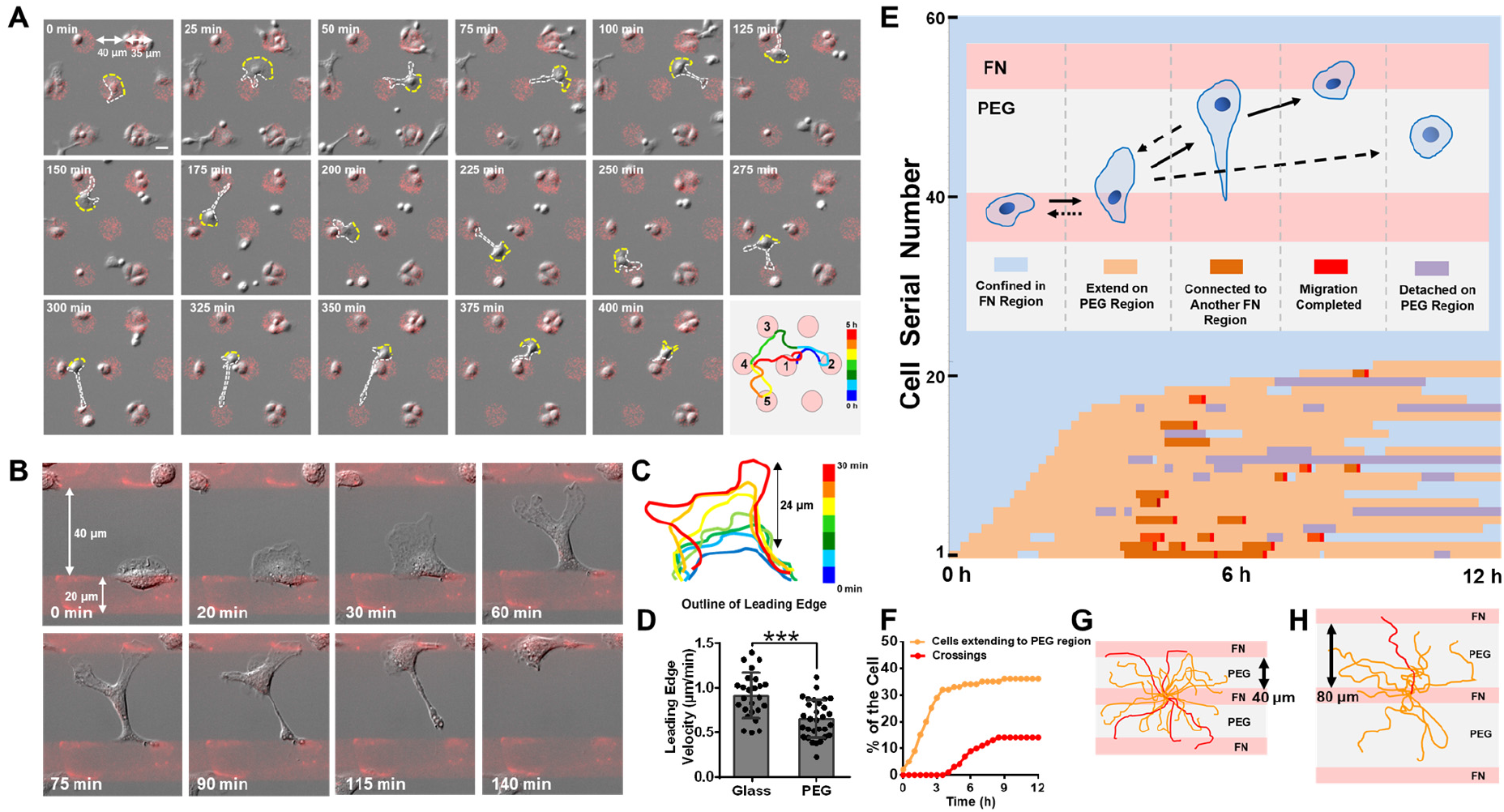
Macrophages migrate across non-adhesive regions to adhesive FN regions in a mesenchymal mode. **(A)** A macrophage migrates from one adhesive FN island (diameter = 35 μm) to another island (gap distance=40 μm) five times in 6 hours. Color indicates different time points. Scale bar=20 μm. **(B)** Macrophage migrates across a striped FN-PEG pattern. The FN stripe is 20 μm in width with a gap distance of 40 μm. **(C)** Outline of the macrophage leading edge in (B). Color indicates different time points. **(D)** Statistical velocity of the leading edge on the glass and PEG surfaces. Statistics were performed by Student’s t test. Asterisks indicate statistical significance compared with the glass group, ***P < 0.001. **(E)** Statistical schematic of macrophage migration on the alternate non-adhesive surface. Watery blue, yellow, orange, red, and gray represented migration states, including cells being confined in the FN region, extending to the PEG region, being elongated and connecting to another FN region, a crossing completed, and being detached on the PEG region, respectively. **(F)** Quantification of the fraction of cells extending to PEG and crossings in (E) over time. Migration trajectory of macrophages on striped patterns with gap distances of 40 μm **(G)** and 80 μm **(H)**. Red traces represent completed crossings. Data was obtained from movie S1, part II.

To quantitatively describe the statistical behavior of macrophages, we employed different colors to define different migration states of macrophages on alternate non-adhesive surfaces. That is, watery blue, yellow, orange, red and violet represented states of cells being confined in the FN region, extending on the PEG region, being elongated and connecting to another FN region, migration across-PEG region completed (we styled that as “a crossing”) and being detached on the PEG region, respectively. Each horizontal line represented one single cell, while different colors at different time points indicated the transition of migration states (Fig.1E). The selected view contained 60 cells, and the time-lapse imaging lasted 12 hours (Movie S1, part III). All cells in the view were lined up according to the time order of extending to the PEG region. It was shown in Fig.1F that the ratio of cells that were able to extend to the PEG region peaked in 3 hours at ∼36%, and that ∼20% of the cell could complete at least one crossing. The trajectory of cells in movie S1, part III indicated random locomotion (only the first crossing was recorded). Varying the gap distance, we found a maximal PEG gap distance of ∼80 μm that macrophage could make a crossing (Fig. 1 H; movie S1, part IV).

We defined this locomotion by two abilities, namely, the ability to extend the cell body to the non-adhesive PEG region and the ability to elongate the cell body to reach another adhesive region. To demonstrate the uniqueness of macrophage motility, we observed the migration of other 25 types of cells on alternate non-adhesive surfaces. Almost none of the examined cancer cells, cell lines, or primary cells exhibited any ability to migrate across the alternate non-adhesive surfaces (Table. S1; movie S2, part I-III). Human neutrophil was observed to protrude out of the FN region in an amoeboid phenotype but could not elongate the cell body to reach another FN region. A few Raw264.7 cells, a mouse macrophage cell line, were observed to extend onto the PEG region (Movie S2, part IV). Only mouse peritoneal macrophage and rat peritoneal macrophage (Movie S2, part IV) had a complete ability to migrate across the non-adhesive PEG region. Thus, we suggest that this motility on alternate non-adhesive surfaces is a unique ability for macrophages.

To test whether this macrophage motility was PEG-dependent, we replaced PEG with another cell-repelling regent, F127. It showed that macrophages could also migrate across F127-based alternate non-adhesive surfaces, proving the universality of this motility on non-adhesive surfaces (Movie S2, part IV).

### Macrophage migration on alternate non-adhesive surfaces needs pre-adhesion on the FN region

To characterize the motility of macrophages on alternate non-adhesive surfaces, we first cultured macrophages on a pure PEG surface. We found that macrophages kept a non-adherent round state (Fig. 2A; movie S3, part I). However, when the cell touched the rim of the FN region, it spread rapidly on the adhesive FN region (Fig. 2B; movie S3, part I). Further, we observed that adhered cells elongated cell body to the PEG region and retracted (Fig. 2C; movie S3, part II), or else they would lose adhesion on the FN region and subsequently turn round-shaped on the PEG region in a dozen minutes (Fig. 2D; movie S3, part II), suggesting that macrophage motility needed a pre-adhesion on the FN region.

**Figure 2.**
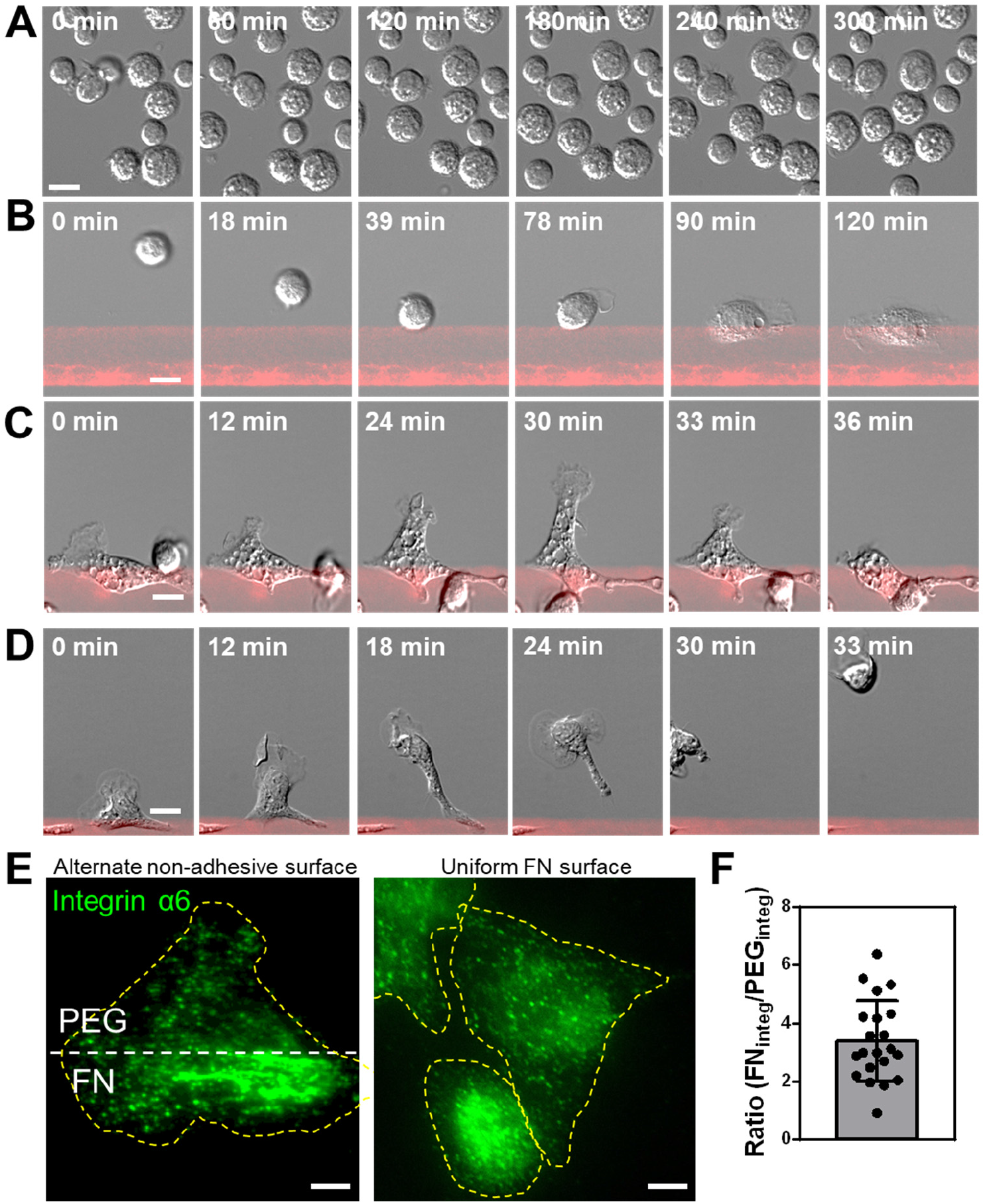
Macrophage migration on non-adhesive surfaces needs pre-adhesive on the FN region. **(A)** Macrophages keep a suspending round morphology on the pure PEG region. **(B)** Macrophage adheres and spreads when it reaches a FN region. **(C)** Macrophage extends out to PEG region and then retracts. **(D)** Macrophage extends out to the PEG region and then detaches from the FN region followed by turning round. Scale bars, 10 μm from (A) to (D). **(E)** Immunofluorescence image of a macrophage labeled with integrin α6 on alternate non-adhesive surface and uniform FN surface. Dashed white line represents the boundary of the PEG and FN regions. Dashed yellow lines represent the outline of the cells. Scale bars, 5 μm. **(F)** Statistical intensity ratio of integrin α6 between the FN and PEG regions.

Cells used integrin to facilitate cell-ECM adhesion. Thus, we examined the distribution of integrin α6 in macrophages on the alternate non-adhesive surfaces. The results showed preferred assembly of integrin α6 on the FN region (Fig 2E; fig. S2A) with an intensity ratio (Intensity_FN_/Intensity_PEG_) of 3.4±1.4 (Fig 2F). By contrast, integrin α6 was homogeneously distributed throughout the cells cultured on uniform FN substrates (Fig 2E; fig. S2B). Therefore, the heterogeneous distribution of integrin α6 was probably associated with pre-adhesion to the FN region for macrophage motility.

### Podosomes support macrophage migration on alternate non-adhesive patterns

It is widely accepted that cells require adhesion to resist liquid fluctuation. Thus, macrophages are likely to form adhesion on the PEG region rather than suspend on the PEG region. Scanning electron microscopy suggested that macrophages adhered tightly to the PEG surface without interspaces (Fig. S3A). Also, gentle mechanical stimulations could not move the cell on the PEG sregion (Fig. S3B; movie S4), suggesting that macrophages indeed adhered onto the non-adhesive PEG region.

To investigate why macrophages could extend and adhere on the non-adhesive surface, we first labeled F-actin and α-tubulin in fixed cells. We thus found that F-actin was highly enriched in the PEG regions in the form of podosomes *(16)*, while microtubules distributed without region-specificity (Fig. 3A). Using other micropatterns, including dot arrays and meshwork patterns, we further showed that podosomes were exclusively distributed on the PEG regions but avoided the FN regions (Fig. 3B; fig. S3C). Other cells with this motility, including Raw264.7 cells and rat macrophages, also formed dense podosomes on the PEG regions (Fig, S4, A and B). Similar results were observed on the F127 surface for mouse macrophages (Fig. S5C), suggesting that the formation of podosomes was not PEG-dependent.

**Figure 3.**
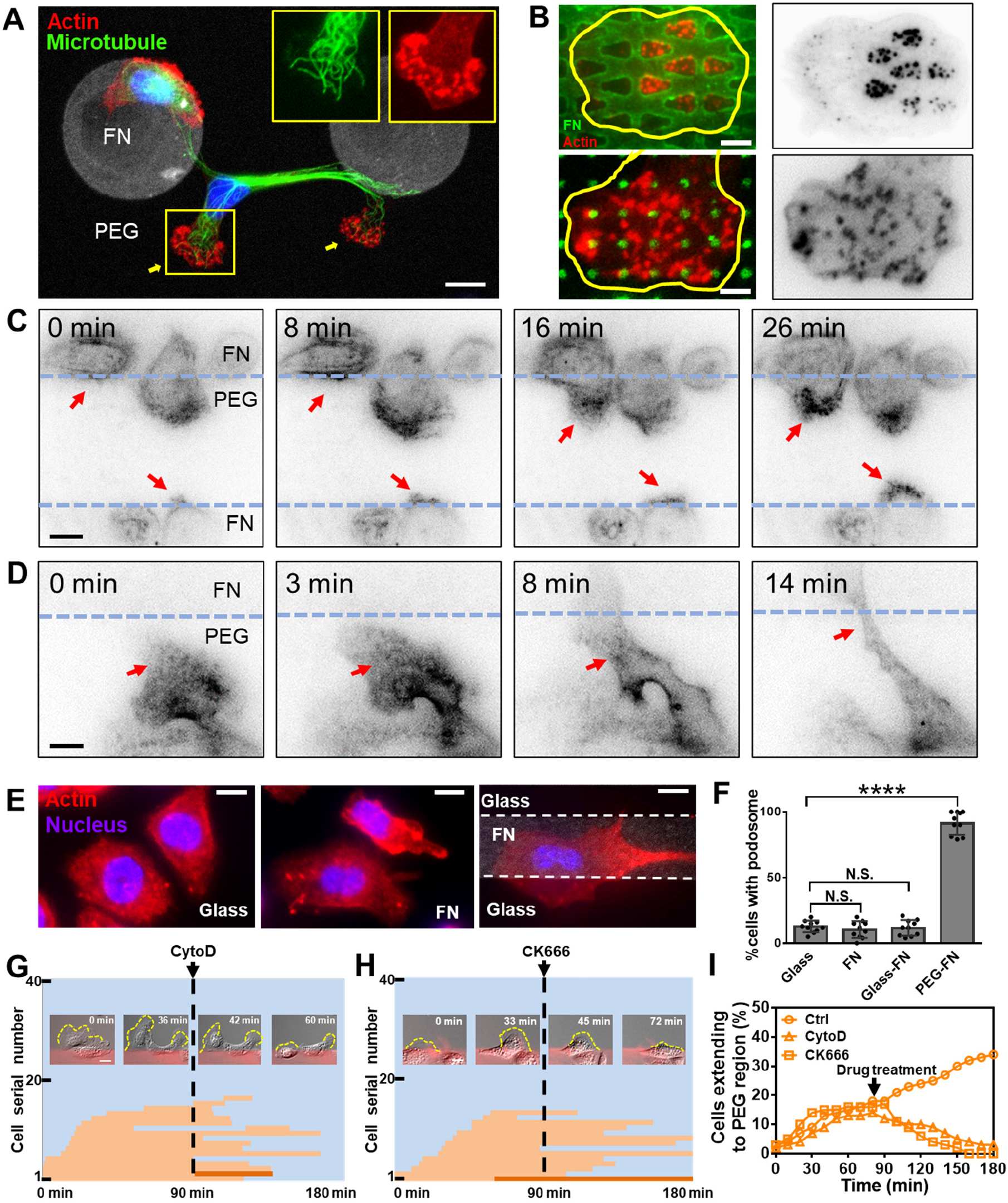
Podosomes facilitate macrophage migration on the alternate non-adhesive region. **(A)** Immunostaining images of macrophage labeled with tubulin (green) and actin (actin). Podosomes were enriched on the PEG region. Scale bar, 10μm. **(B)** Macrophage on meshwork pattern and dots-array pattern labeled by phalloidin. Podosomes avoided the FN regions. Scale bars, 10 μm. **(C)** Macrophage forms podosomes when extending to the PEG region (labeled by SiR-actin). Scale bar, 5 μm. **(D)** Podosomes disassemble when this part reaches the FN region. Scale bar, 5 μm. **(E)** Macrophages do not form podosomes on glass, FN, or glass-FN surface, respectively. Scale bars, 10 μm. **(F)** Quantification of the ratio of podosome-positive cells cultured on glass, FN, FN-glass, and PEG-FN. Statistics were performed by Student’s t test. Asterisks indicate statistical significance compared with the glass group, N.S., no significance, ***P < 0.001. macrophage migration diagram and timelapse-imaging of CytoD **(G)** and **(H)** CK666 group. Scale bars, 20 μm. **(I)** Fraction of cells extending to the PEG region over time in the CytoD and CK666 groups.

Using SiR-actin, a live-cell F-actin labeling probe, we next examined the dynamic of podosomes in macrophages using TIRF microscopy. Podosomes clearly appeared when macrophages extended to the PEG region (Fig. 3C; movie S5, part I) and disappeared fast when they reached another FN region (Fig. 3D; movie S5, part II) or retracted back to the FN region (movie S5, part III). Furthermore, cells on glass, FN, and glass-FN substrates showed no evident podosomes (Fig. 3E). Statistical data showed that the percentages of podosome-positive macrophages were 12.8±1.4%, 10.8±2.0%, 11.7±1.9% and 91.4±2.8% on the glass, FN, glass-FN and PEG-FN substrates, respectively (Fig. 3F). Inhibiting podosome formation by cytochalasin D (CytoD, an actin polymerization inhibitor) or CK666 (an Arp2/3 inhibitor) decreased macrophage motility on the alternative non-adhesive substrates, suggesting a supporting role of podosomes (Fig. 3, G to I; fig. S5; and movie S6).

We also observed dynamic protuberances in the thin lamellipodium layer in DIC images (Movie S7). Fluorescence-labeled F-actin revealed that the protuberances in DIC images corresponded to podosomes (Fig. S6A). Interestingly, we observed membrane protruded after the appearance of a single proximal podosome on the PEG region (Fig. S6B; movie S7), suggesting that podosomes drove the actin polymerization and membrane protrusion on the PEG surfaces.

### Inhibition of Myosin IIA promotes macrophage motility on alternate non-adhesive surfaces

The cytoskeleton determines the mechanical property and migration state of the cell. Hence, we reasoned that adhesion-associated proteins might play a role in macrophage motility. Applying 3D-STORM *(17)*, we revealed the axial organization of actin, paxillin, vinculin, and myosin IIA. Actin on the PEG regions localized ∼50 nm higher than that on the FN regions because of highly-enriched podosomes on the PEG regions (Fig. S7A). Paxillin and vinculin on the FN and PEG surfaces were similar in height (Fig. S7, B and C). Myosin IIA, however, was found to localize ∼100 nm higher on the PEG surface than that on the FN surface in the same cell (Fig. 4, A and B; fig. S7D; and fig.S8C). To exclude the possibility that the PEG surface was physically higher than the FN surface, we performed atom force microscopy (AFM) to detect the surface topography of alternate non-adhesive micropatterns. Results showed that the height of micropatterned FN region was ∼10 nm higher than the PEG region (Fig.S7E), below the resolution limit of 3D-STORM. Statistical height differences (Δh=***Peak***_PEG_-***Peak***_FN_) were 53.7±6.5 nm, 25.9±4.7 nm, 25.8±6.4 nm, and 105.2±7.6 nm for actin, paxillin, vinculin, and myosin, respectively (Fig. S7F). A conclusive schematic diagram of protein distribution of macrophages extending to the PEG region is displayed in Fig. S7G.

**Figure 4.**
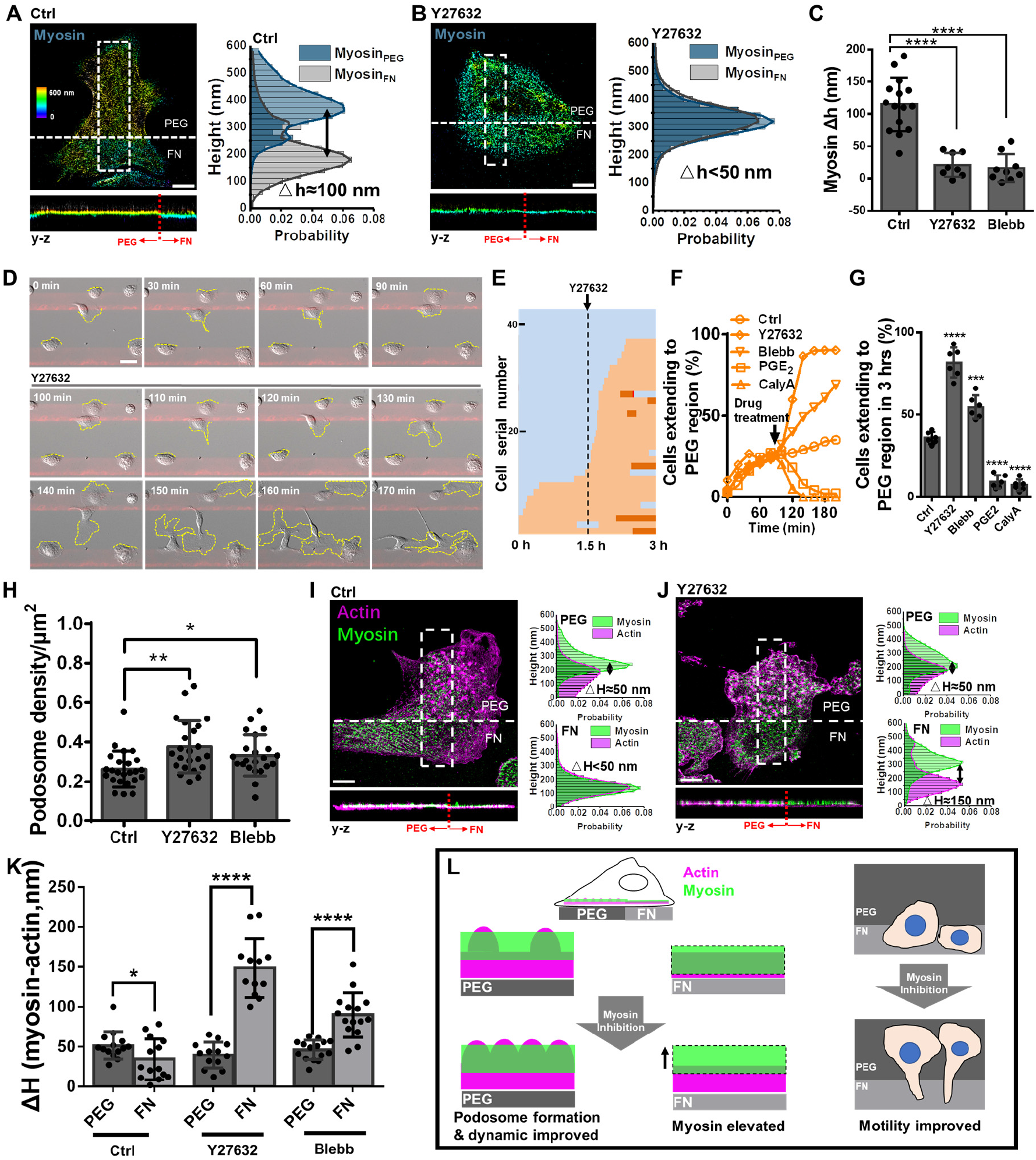
Effect of myosin IIA on macrophage motility on the alternate non-adhesive PEG surface. **(A and B)** Representative 3D-STORM images of myosin in individual macrophages at the PEG-FN boundary of the ctrl (A) or Y27632 (B) groups. The z-position is color-coded according to the color bar, with violet indicating positions closest to the substratum and red indicating farthest. The white boxes are selected regions, and their corresponding zoom-in images of z direction are shown below. The z profiles of myosin on the PEG and FN regions along the vertical section in the box is shown on the right. Scale bar, 5 μm. **(C)** Quantification of myosin IIA △h in the ctrl, Y27643, and Blebb groups. Myosin△h=**Peak**_PEG_-**Peak**_FN_. **(D)** Y27632 promotes macrophage motility on the alternate non-adhesive PEG surface. Scale bar, 10 μm. **(E)** Macrophage migration diagram of Y27632 group. **(F)** Fraction of cells extending to the PEG region over time in the ctrl, Y27632, Blebb, PGE_2,_ and CalyA groups. **(G)** Quantification of the ratio of cells extending to the PEG region in different groups 1.5 hrs after the different drug treatments. **(H)** Quantification of podosome density in the ctrl, Y27632, and Blebb groups. **(I, J)** Representative two-color 3D-STORM images of actin (magenta) and myosin (green) in individual macrophages at the PEG-FN boundary for the ctrl and Y27632 groups. The white boxes are selected regions, and the corresponding zoom-in images of z direction are shown below. The z profiles of actin and myosin on the PEG and FN regions along the vertical section in the box are shown on the right. Scale bars, 5 μm. **(K)** Quantification of myosin△H in ctrl, Y27643, and Blebb groups on the PEG and FN regions. △H=**Peak**_myosin_-**Peak**_actin_. **(L)** Inhibition of myosin II promotes podosome density and macrophage motility on the PEG region.

To further gain insights into the role of myosin IIA in macrophage motility, we applied Y27632 (a ROCK inhibitor) and Blebbistatin (Blebb, a myosin II ATPase inhibitor). Results showed that these drugs eliminated the height difference of Myosin IIA in the PEG and FN regions (Fig. 4, A to C; fig. S8, D and E). Statistical Δh of myosin IIA was 109.2±14.9 nm, 18.3±7.6 nm, and 12.0±9.4 nm for the control group, Y27632 group, and Blebb group, respectively (Fig. 4C). Furthermore, the macrophage motility on the alternate non-adhesive substrates was enhanced greatly upon myosin IIA inhibition (Fig. 4, D to G; fig. S8, A and B; and movie. S8), while activating myosin IIA pathway by Prostaglandin E_2_ (PGE_2_, a Rho-ROCK pathway activator) and Calyculin A (CalyA, a phosphatase inhibitor) was found to reduce the macrophage motility (Fig. 4, F and G; fig. S9, A and B; and movie. S8). In detail, the ratio of cells extending to the PEG region within 3 hours was 35.9±1.9%, 81.7±3.7%, 54.7±2.9%, 7.2±2.0%, and 6.1±1.4% for the control, Y27632, Blebb, PGE_2_ and CalyA groups, respectively (Fig. 4G). Inhibition of myosin IIA also increased the density of podosomes, which was 0.26±0.01/μm^2^, 0.37±0.03/μm^2^, and 0.33±0.02/μm^2^ for the control, Y27632, and Blebb groups (Fig. 4H). By contrast, activating myosin IIA by PGE_2_ and CalyA blocked the formation of podosomes (Fig. S9, C and D). Therefore, activation of the Rho-ROCK-myosin IIA pathway played a negative role in podosome formation and macrophage motility on the alternate non-adhesive surface.

Spatial coordination of myosin and actin at the cortex regulates cell surface mechanics *(18)*. Hence, we analyzed the relative localization of cortical actin and myosin IIA in macrophages on the alternate non-adhesive surface. It was observed that in the control group, actin and myosin almost overlapped on the FN region and on normal substrates, while myosin IIA was ∼50 nm higher than actin on the PEG region (Fig.4I; fig. S10, A and B). Notably, when myosin IIA was inhibited by Blebb or Y27632, myosin IIA localized higher towards the cytoplasm and separated from the actin cortex on the FN region (Fig. 4J; fig. S10, C and D), implicating a reduction of cortex tension as previously reported *(17)*. The statistical height differences of actin and myosin II ΔH (ΔH=***Peak***_myosin_-***Peak***_actin_) were 51.4±4.6 nm, 31.8±4.9 nm, 45.9±3.7 nm on the PEG region for the control, Y27632, and Blebb groups. ΔH were 46.5±4.6 nm, 148.6±10.7 nm, and 89.6±7.1 nm on the FN region for the control, Y27632, and Blebb groups (Fig. 4K). Therefore, the reduction of cortex tension may promote macrophage motility on the alternate non-adhesive surface (Fig. 4L).

### A developed CPM model for macrophage motility on alternate non-adhesive surfaces

The above intriguing observations of macrophages led us to develop a cellular Potts model (CPM)-based theoretical framework to describe the motility of macrophages on the alternate non-adhesive surface *(19)*. In this model, cells possessed an intrinsic activity gradient of actin that led to migration *(20, 21)*. Podosomes, which enable adhesion and locomotion on PEG, were induced by the PEG surface when cells reached the PEG-FN boundary. A positive feedback path was constructed that podosomes supported the actin polymerization and protrusion (Fig. S7B; movie. S7), while the activating of actin promoted the formation of podosomes (Fig. 5A). A negative pathway was set that podosomes disassembled when cells reached another FN region. Detailed equations and parameters were described in Materials and methods. The simulation results showed that actin polarization produced a leading edge of the cell with podosomes appearing when extending to the PEG region (Fig. 5B; movie. S9, part I). When the uropod of the cell was detached from FN, the cell turned round on the PEG region in a dozen minutes (Fig. 5C; movie. S9, part II). Afterwards, the cell reached another FN region and re-adhered to it (Fig. 5C; movie. S9, part II). Considering that inhibition of myosin II increased the density of podosomes (Fig. 4H), we up-regulated the probability of podosome generation and resulted in a higher probability of crossing as a consequence (Fig. 5, D and E; movie. S9, part III). Moreover, macrophage migration in a triangular lattice pattern was also simulated in movie. S9, part IV. These simulation results were highly consistent with our experimental observations (Fig. 1, A and B; fig. 2B; and fig. 4, D to G).

**Figure 5.**
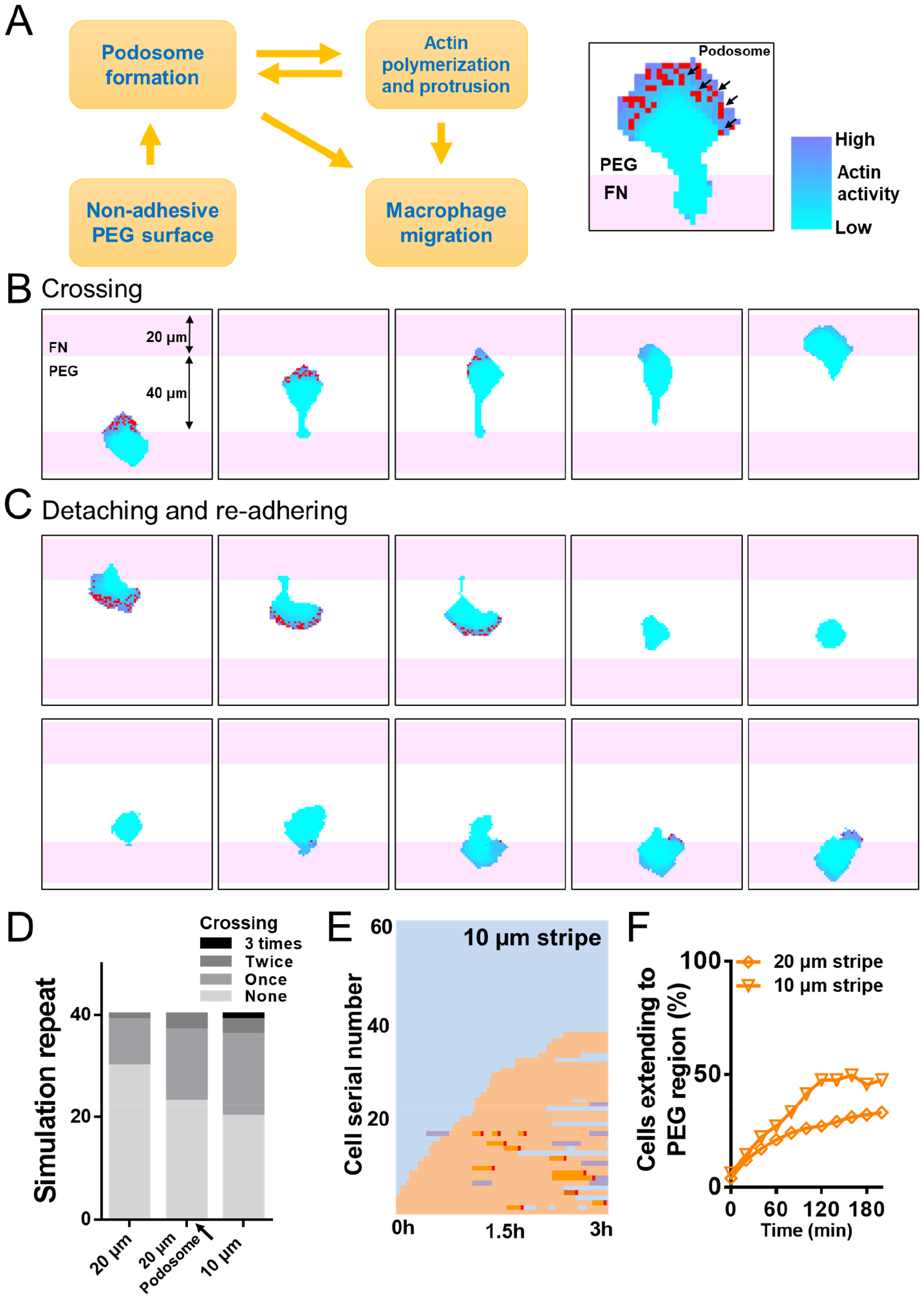
Theoretical simulation of macrophage motility on the alternate non-adhesive surface. **(A)** Schematic of the relationship between podosome formation, actin polymerization, PEG and macrophage migration in the developed CPM model. Color bar encodes the intensity of actin activity. Red pixels indicate the podosomes. **(B)** Simulation presentation of a macrophage crossing a 40 μm PEG gap. **(C)** Simulation presentation of macrophage detaching, turning round, and re-adhering on the alternate non-adhesive surface. **(D)** Number of crossings in 40 repeats of simulation in 20 μm FN stripe, 20 μm FN stripe with podosome up-regulated and 10 μm FN stripe groups. **(E)** Macrophage migration diagram of 10 μm FN stripe group from experimental results of movie S10. Cells could extend to the PEG region and complete crossings on the 10 μm FN stripe more easily than cells on the 20 μm FN stripe. **(F)** Fraction of cells extending to the PEG region over time in the 20 μm and 10 μm FN stripe groups.

To further validate our model, we simulated the cell migration on narrow FN stripes of 10 μm. It was found that the cell extended to the PEG region more easily than on wider 20 μm stripes with a higher efficiency of crossing (Movie. S10, part I). Subsequent experimental results showed that narrow stripes of 10 μm indeed enhanced the macrophage motility (Fig. 5, E and F; movie. S10, part II). Together, these results demonstrated that our model showed good agreement with experimental results as well as made predictions to guide further experiments.

## Discussion

Cell migration is essential for varieties of biological processes. It is widely acknowledged that mesenchymal cells cannot migrate on non-adhesive substrates lacking adhesion to generate force. Recently, it was reported that slow mesenchymal cells *(e*.*g*., NHDF cells, Hela cells and MDCK cells) under physical confinement could convert to amoeboid migration on non-adhesive surfaces *(22)*. In the present work, we found that mesenchymal migration can happen on alternate non-adhesive surfaces in macrophages.

*In vivo*, immune cells need to perform effective migration through complex ECM gaps unsuitable for adhesion (e.g., collagen-free zones) in response to infection or injury *(9,23-25)*. This migration through the ECM gaps was mostly associated with the amoeboid mode *(23)*, such as a swimming-like locomotion of neutrophil *(26,27)*. Recent studies also observed that some cells, including NIH 3T3 cells *(28)* and human umbilical cord vascular endothelial cells *(29)*, could overcome short gaps of ∼10 μm *in vitro*. Of particular note is that those two cells used protrusions to move across gaps much smaller than the cell size without extreme elongation of the cell body or adhesion on non-adhesive surfaces. Quite differently, macrophages could migrate across alternate non-adhesive PEG gap as wide as 80 μm in a mesenchymal mode. This new behavior of migration endows macrophages with enormous adaptability to migrate through complex interstitial tissues.

Furthermore, our results showed that macrophages could not adhere on a pure PEG surface but needed pre-adhesion on the FN region for further locomotion (Fig. 2, A to D). Only when the cell reached another FN region would it dissociate the uropod and accomplish migration across the PEG gap (Fig. 1, A and B). These results indicated a whole-cell back-front-back signaling pathway that regulated the adhesion and migration of macrophages on the alternate non-adhesive surface. In addition, podosomes were found to largely appear only on the PEG region and disappeared quickly when this part reached an FN region (Fig. 3, C and D; movie S5). Therefore, the formation of podosomes needed a prior activation of adhesion signaling, but was mutually exclusive in distribution with the adhesive FN region.

We tested 25 other types of cells on alternate non-adhesive surfaces. Only macrophage-related cells exhibited such motility, including mouse macrophage, rat macrophage, and raw264.7 cells (Table. S1). This unique capacity is probably explained by podosomes according to our results (Fig. 3), which facilitated the adhesion and migration on the PEG surface (Fig. S6; movie S7). Previous studies reported that podosomes and other adhesion structures assembled on the FN adhesive region *(30-32)*, which was distinguished from our results. We reason that different types of cells might exhibit diverse forms and functions of podosomes (i.e., a rosette-shaped podosome ring of microglia *(33)* and invadopodia of cancer cells *(34)*), leading to a different spatial distribution of podosomes and distinct motility on the alternate non-adhesive surface. Besides, our simulation model built on the supporting role of podosomes (Fig. 5A; movie S9) reproduced our experimental observations, confirming the validity of this assumption from both experimental and theoretical perspectives (Movie S9; movie S10).

Previous studies have reported different effects of myosin IIA on podosome formation, including promotional *(35)*, inhibitory *(36)*, and null effects *(37,38)*. Our results suggested that myosin IIA played a negative role in the formation of podosomes in macrophages. Inhibition of myosin IIA greatly increased the motility of macrophages on the alternate non-adhesive surface via facilitating podosome formation (Fig. 4, D to G) from the perspective of the signaling pathway (Fig. S11). In our mathematical model, up-regulation of podosome generation simulated the effect of myosin IIA inhibition, which led to an enhancement of macrophage motility (Movie S9, part III). On the other hand, myosin IIA could provide the intracellular contraction force *(39)*, which could impede cell extension to the PEG region. Inhibition of myosin IIA countered the contraction force in the cortex and thus promoted cell extension to the PEG region from the perspective of cell surface mechanism (Fig. 4, I to K). Therefore, counteracting membrane tension may be another approach to improve macrophage motility on the alternative non-adhesive surface.

Overall, our work uncovered a new migratory behavior of macrophages on alternate non-adhesive surfaces, which gain new insight into macrophage motility, as well as enrich our understanding of cell migration.

## Acknowledgments

We thank Prof. Xueliang Zhu for the helpful discussions.

## Author contributions

L. P. conceived the research and were in charge of the overall direction. L. P. and F. X. designed the experiments. F. X. and P. Z. performed the experiments. F. X. and H. D. performed computational modeling. F. X., J. Y., C.F., M. H. and F.H. analyzed the data. J. Z. and L.C. contributed to the interpretation of the results. F. X. and L. P. wrote the manuscript. L. P. and J. X. supervised the work.

## Data and materials availability

The codes for the CPM model are available at https://github.com/Donnie3246/Xing_et_al_2022.git.

## Supplementary Materials

Materials and Methods

Supplementary Text

Figs. S1 to S11

Tables S1

References *(38*–*45)*

Movies S1 to S10

## References

1. T. A. Wynn, A. Chawla, J. W. Pollard, Macrophage biology in development, homeostasis and disease. Nature 496, 445–455 (2013).

2. P. J. Murray, Macrophage polarization. Annu. Rev. Physiol. 79, 541–566 (2017).

3. M. H. Raymond et al., Live cell tracking of macrophage efferocytosis during Drosophila embryo development in vivo. Science 375, 1182–1187 (2022).

4. D. S. Eom, D. M. Parichy, A macrophage relay for long-distance signaling during postembryonic tissue remodeling. Science 355, 1317–1320 (2017).

5. L. C. Davies, S. J. Jenkins, J. E. Allen, R. T. Philip, Tissue-resident macrophages. Nat. Immunol. 14, 986–995 (2013).

6. B. Stolp et al., Salivary gland macrophages and tissue-resident CD8+ T cells cooperate for homeostatic organ surveillance. Sci. Immunol. 5, eaaz4371 (2020).

7. C. V. Jakubzick, G. J. Randolph, P. M. Henson. Monocyte differentiation and antigen-presenting functions. Nat. Rev. Immunol. 17, 349–362 (2017).

8. T. Gerhardt, K. Ley. Monocyte trafficking across the vessel wall. Cardiovasc. Res. 107, 321–330 (2015).

9. J. L. Orgaz, et al., Diverse matrix metalloproteinase functions regulate cancer amoeboid migration. Nat. Commun. 5, 1–13 (2014)

10. F. Barros-Becker et al., Distinct tissue damage and microbial cues drive neutrophil and macrophage recruitment to thermal injury. Iscience 23, 101699 (2020).

11. F. Barros-Becker, P. Y. Lam, R. Fisher, A. Huttenlocher, Live imaging reveals distinct modes of neutrophil and macrophage migration within interstitial tissues. J. Cell Sci. 130, 3801–3808 (2017).

12. A. J. Davidson, W. Wood. Macrophages use distinct actin regulators to switch engulfment strategies and ensure phagocytic plasticity in vivo. Cell Rep. 31, 107692 (2020).

13. R. Sridharan, B. Cavanagh, A. R. Cameron, D. J. Kelly, F. J. O’Brien, Material stiffness influences the polarization state, function and migration mode of macrophages. Acta Biomater. 89, 47–59 (2019).

14. F. Y. McWhorter, T. Wang, P. Nguyen, W. F. Liu, Modulation of macrophage phenotype by cell shape. Proc. Natl. Acad. Sci. U.S.A. 110, 17253-17258 (13).

15. K.M. Hansson, et al. Whole blood coagulation on protein adsorption-resistant PEG and peptide functionalized PEG-coated titanium surfaces. Biomaterials 26, 861–872 (2005).

16. H. Schachtner, S. D. J. Calaminus, S. G. Thomas et al. Podosomes in adhesion, migration, mechanosensing and matrix remodeling. Cytoskeleton 70, 572–589 (2013).

17. B. Huang, X. Wang, M. Bates, X. Zhuang. Three-dimensional super-resolution imaging by stochastic optical reconstruction microscopy. Science 319, 810–813 (2008).

18. B. A. Truong Quang et al., Extent of myosin penetration within the actin cortex regulates cell surface mechanics. Nat. Commun. 12, 1–12 (2021).

19. F. Graner, J. A. Glazier. Simulation of biological cell sorting using a two-dimensional extended Potts model. Phys. Rev. Lett. 69, 2013 (1992).

20. I. Niculescu, J. Textor, R. J. De Boer. Crawling and gliding: a computational model for shape-driven cell migration. PLoS Comput. Biol. 11, e1004280 (2015).

21. I. M. N. Wortel et al. Local actin dynamics couple speed and persistence in a cellular Potts model of cell migration. Biophys. J. 120, 2609–2622 (2021).

22. Y. J. Liu et al., Confinement and low adhesion induce fast amoeboid migration of slow mesenchymal cells. Cell 160, 659–672 (2015).

23. P. Friedl, B. Weigelin, Interstitial leukocyte migration and immune function. Nat. Immunol. 9, 960–969 (2008).

24. M. Krause, K. Wolf, Cancer cell migration in 3D tissue: Negotiating space by proteolysis and nuclear deformability. Cell Adhes. Migr. 9, 357–366 (2015).

25. K. M. Yamada, M. Sixt, Mechanisms of 3D cell migration. Nat. Rev. Mol. Cell Biol. 20, 738–752 (2019).

26. N. P. Barry, M. S. Bretscher. Dictyostelium amoebae and neutrophils can swim. Proc. Natl. Acad. Sci. U.S.A. 107, 11376–11380 (2010).

27. L. Aoun et al., Amoeboid swimming is propelled by molecular paddling in lymphocytes. Biophys. J. 119, 1157–1177 (2020).

28. S. L. Vecchio et al., Collective dynamics of focal adhesions regulate direction of cell motion. Cell Syst. 10, 535–542 (2020).

29. D. Garbett et al., T-Plastin reinforces membrane protrusions to bridge matrix gaps during cell migration. Nat. Commun. 11, 1–18 (2020).

30. A. Labernadie, C. Thibault, C. Vieu, I. Maridonneau-Parini, G. M. Charrière, Dynamics of podosome stiffness revealed by atomic force microscopy. Proc. Natl. Acad. Sci. U.S.A. 107, 21016–21021 (2010).

31. C. Yu et al., Integrin-matrix clusters form podosome-like adhesions in the absence of traction forces. Cell Rep. 5, 1456–1468 (2013).

32. N. B. M. Rafiq et al., A mechano-signalling network linking microtubules, myosin IIA filaments and integrin-based adhesions. Nat. Mater. 18, 638–649 (2019).

33. C. Vincent, T. A. Siddiqui, L. C. Schlichter. Podosomes in migrating microglia: components and matrix degradation. J. Neuroinflamm. 9, 1–15 (2012).

34. E. Dalaka et al., Direct measurement of vertical forces shows correlation between mechanical activity and proteolytic ability of invadopodia. Sci. Adv. 6, eaax6912 (2020).

35. H. Schachtner et al., Megakaryocytes assemble podosomes that degrade matrix and protrude through basement membrane. Blood 121, 2542–2552 (2013).

36. O. Collin et al. Self-organized podosomes are dynamic mechanosensors. Curr. Biol. 18, 1288–1294 (2008).

37. K. Van Den Dries et al., Interplay between myosin IIA-mediated contractility and actin network integrity orchestrates podosome composition and oscillations. Nat. Commun. 4, 1–13 (2013).

38. S. F. G. van Helden et al. PGE_2_-mediated podosome loss in dendritic cells is dependent on actomyosin contraction downstream of the RhoA–Rho-kinase axis. J. Cell Sci. 121, 1096–1106 (2008).

39. O. Milberg et al., Concerted actions of distinct nonmuscle myosin II isoforms drive intracellular membrane remodeling in live animals. J. Cell Biol. 216, 1925–1936 (2017).

40. L. Pan, X. Zhang, K. Song, J. Xu. Exogenous nitric oxide-induced release of calcium from intracellular IP3 receptor-sensitive stores via S-nitrosylation in respiratory burst-dependent neutrophils. Biochem. Biophys. Res. Commun. 377, 1320–1325 (2008).

41. M. Pannell et al. Imaging of translocator protein upregulation is selective for pro-inflammatory polarized astrocytes and microglia. Glia 68, 280–297 (2020).

42. H. Ghasemi Hamidabadi et al. Chitosan-intercalated montmorillonite/poly (vinyl alcohol) nanofibers as a platform to guide neuronlike differentiation of human dental pulp stem cells. ACS Appl. Mater. Interfaces 9, 11392–11404 (2017).

43. S. Zhu et al. Involvement of transient receptor potential melastatin-8 (TRPM8) in menthol-induced calcium entry, reactive oxygen species production and cell death in rheumatoid arthritis rat synovial fibroblasts. Eur. J. Pharmacol. 725, 1–9 (2014).

44. T. H. Chen et al. Directing tissue morphogenesis via self-assembly of vascular mesenchymal cells. Biomaterials 33, 9019–9026 (2012).

45. F. Xing et al. Spatiotemporal characteristics of intercellular calcium wave communication in micropatterned assemblies of single cells. ACS Appl. Mater. Interfaces 10, 2937–2945 (2018).

